# Top-down Control of Inhibition Reshapes Neural Dynamics Giving Rise to a Diversity of Computations

**DOI:** 10.1101/2020.02.25.964965

**Authors:** Zhen Chen, Krishnan Padmanabhan

## Abstract

Growing evidence shows that top-down projections from excitatory neurons in higher brain areas selectively synapse onto local inhibitory interneurons in sensory systems. While this connectivity is conserved across sensory modalities, the role of this feedback in shaping the dynamics of local circuits, and the resultant computational benefits it provides remains poorly understood. Using rate models of neuronal firing in a network consisting of excitatory, inhibitory and top-down populations, we found that changes in the weight of feedback to inhibitory neurons generated diverse network dynamics and complex transitions between these dynamics. Additionally, modulation of the weight of top-down feedback supported a number of computations, including both pattern separation and oscillatory synchrony. A bifurcation analysis of the network identified a new mechanism by which gamma oscillations could be generated in a model of neural circuits, which we termed **T**op-down control of **I**nhibitory **N**euron **G**amma (TING). We identified the unique roles that top-down feedback of inhibition plays in shaping network dynamics and computation, and the ways in which these dynamics can be deployed to process sensory inputs.

**Significance Statement:** The functional role of feedback projections, connecting excitatory neurons in higher brain areas to inhibitory neurons in primary sensory regions, remains a fundamental open question in neuroscience. Growing evidence suggests that this architecture is recapitulated across a diverse array of sensory systems, ranging from vision to olfaction. Using a rate model of top-down feedback onto inhibition, we found that changes in the weight of feedback support both pattern separation and oscillatory synchrony, including a mechanism by which top-down inputs could entrain gamma oscillations within local networks. These dual functions were accomplished via a codimension-2 bifurcation in the dynamical system. Our results highlight a key role for this top-down feedback, gating inhibition to facilitate often diametrically different local computations.

## Introduction

While inhibitory interneurons constitute a comparatively small fraction (20%) of neurons in the neocortex (Meinecke and Peters 1987), they encompass a diverse array of types, distinct in gene expression, morphology, and connectivity (Kepecs and Fishell 2014). Thought to be largely local in their influence (Yoshimura and Callaway 2005, Fino and Yuste 2011), the interneurons nonetheless play critical roles in shaping the dynamics of neural circuits (Fishell and Kepecs 2019) across all sensory and motor systems (Urban and Sakmann 2002, Okun and Lampl 2008, Atallah and Scanziani 2009, Cardin, Carlén et al. 2009, Poo and Isaacson 2009, Sohal, Zhang et al. 2009) (Silberberg and Markram 2007, Packer and Yuste 2011). (Buzsáki 1984).

Interestingly, recent anatomical evidence has identified top-down centrifugal afferents from higher cortical areas that specifically target inhibitory interneurons. In visual system of the mouse for example, axons from the cingulate (cG) of the frontal cortex project to GABAerigic inhibitory neurons in V1, and activation of cG neurons improved performance on a visual discrimination task (Zhang, Xu et al. 2014). Similarly, in the olfactory system, axons from excitatory neurons in the piriform cortex synapse onto the local inhibitory granule cells in olfactory bulbs, whereby they can modulate the function of the mitral and tufted cells, the principal relays of olfactory information from the bulb to the brain (Shipley and Adamek 1984, Padmanabhan, Osakada et al. 2016, Padmanabhan, Osakada et al. 2019). These top-down higher area neurons receive input from only excitatory populations in sensory areas, but they exert their influence on the local circuit dynamics via synapses onto inhibitory populations. Consequently, whereas the information relayed to these top-down cells comes from the local excitatory population, they intervene in network dynamics through the inhibitory population. Although this architecture is recapitulated across multiple sensory systems, the computational role of this top-down control of inhibition and the dynamic mechanisms that give rise to such computations remain largely unknown.

To address this question, we built a three-node network model consisting of an excitatory (E), inhibitory (I) and top-down (T) population, and studied how firing rate dynamics were influenced by the structure of the top-down connection weight onto inhibition. Modulating the weight of the top-down connections to local inhibitory neurons reshaped the dynamics of the local excitatory-inhibitory (E-I) circuit in a way that enhanced sensory discrimination as well as generated oscillatory synchrony including entraining gamma oscillations in the local circuit (**T**op-down control of **I**nhibitory **N**euron **G**amma, (TING)). Finally, the mechanism underlying the dynamics, as well as the functional roles played by top-down control of inhibition occurred via a codimension-2 bifurcation in the dynamical system. By gating the magnitude of top-down feedback, a number of seemingly disparate computations could be supported by a single circuit, providing an additional framework for the diversity of inhibitory interneuron function.

## Materials and Methods

### Network Model

The network model was composed of three nodes, a local excitatory (E) and inhibitory (I) nodes which were reciprocally coupled, and a top-down node (T) that received input from the local E node and projected back to the inhibitory I node (Fig.1A). In the model, *r_i_*(*t*),*i* = 1,2,3 represented the firing rates of the three neuron populations respectively, whose dynamics were determined by Wilson-Cowan equations (Wilson and Cowan 1972) as follows:

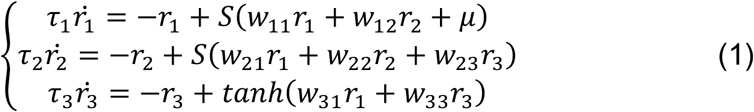

where *S* is the sigmoid function:

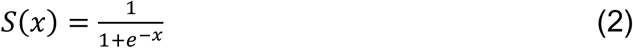

which described the nonlinear relationship between the mean synaptic input and average firing rate (normalized to a range between 0 and 1). The parameter *τ_i_, i* = 1,2,3 was the time constant for each population, characterizing how quickly the dynamics of each population evolved. The excitatory neuron population (E) received an external stimulus *μ*, that represented the only external input to the system (Fig. S1). The connection weight from population *j* to population *i* was denoted by *w_ij_,i,j* = 1,2,3, among which *w*_11_, *w*_21_, *w*_61_, *w*_26_, *w*_66_ > 0 and *w*_12_, *w*_22_ < 0. The connection weight *w_ij_,i,j* = 1,2,3 represented the average synaptic input received by the neuron population i from the population j. Throughout this paper, we set the parameters as follows: *w*_11_ = 8.7, *w*_12_ = −10,*w*_21_ = 7.0, *w*_22_ = −13,*w*_31_ = 1.5, *w*_33_ = 0.5. As there was no inhibitory synaptic input into the top-down population (T), a hyperbolic tangent function *tanh*(*x*) was used for *r*_3_(*t*) to cover the full range of firing rates [0,1] (Fig.1C).

**Figure 1.**
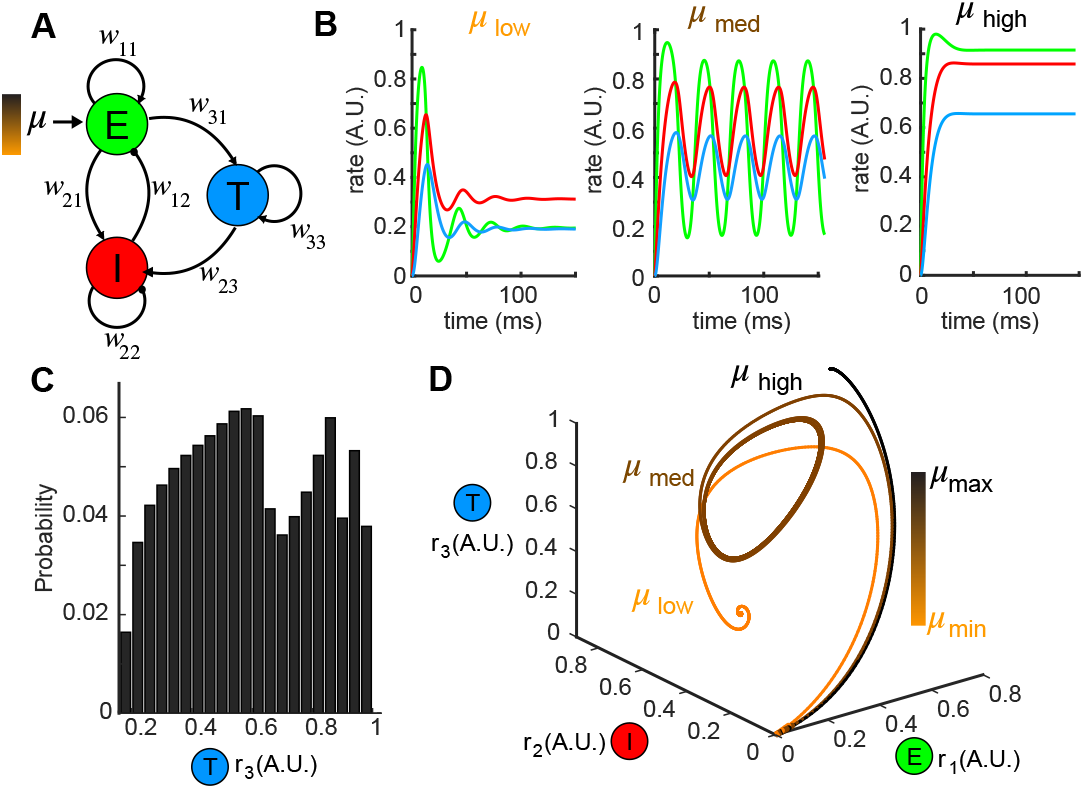
Network model of top-down feedback to inhibitory neurons exhibiting complex dynamics. (A). A schematic diagram illustrating the topology of the network model. Each node denotes a neuron population and the connection weights are defined by *w_ij_* (excitatory: arrows; inhibitory: circles). (B). Network responses (color-coded firing rates match population nodes in (A)) to different stimulus magnitudes (left: *μ*_low_ = 0.1, middle: *μ*_med_ = 1.5, right: *μ*_high_ = 3.0). (C). Distribution of all possible steady states r_3_ shows the diversity of responses the network can generate. (D). Trajectories of the firing rate responses plotted in (B) visualized in the phase space spanned by (*r*_1_*r*_2_, *r*_3_). The color bar indicates stimulus for three representative levels as in (B) (*μ*_low_ = 0.1, *μ*_med_ = 1.5, *μ*_high_ = 3.0).

### Metric definition

As representations of the network (*ω*-limit sets) could take on different forms, an equilibrium or a limit cycle, we defined two quantitative metrics: *d^E^*(Ω_1_,Ω_2_) and *d^S^*(*r*_3_ |_Ω_1__, *r*_3_ |_Ω_2__) which served to measure the distance between different types of *ω*-limit sets in responses to any given stimulus pair (*μ*_1_, *μ*_2_) where *μ*_2_ = *μ*_1_ + Δ. *d^E^*(Ω_1_,Ω_2_) denoted the average Euclidean distance between Ω_1_ and Ω_2_ in the three-dimensional (3D) phase space of firing rates (*r*_1_ *r*_2_, *r*_3_), and the spectrum distance *d^S^*(*r*_3_ |_Ω_1__, *r*_3_ |_Ω_2__) was a sum of the squared differences between both direct components (DC) and alternating components (AC) in the amplitude-frequency domain of the Fourier transforms to the signals *r*_3_(*t*) associated with the two stimuli (Fig. S2-S3, Supplementary material). When both Ω_1_ and Ω_2_ were equilibria, the spectrum distance *d^s^* was a projection of the Euclidean distance *d^E^* onto the *r*_3_ axis. The spectrum distance was however more sensitive to bifurcations when one of the two *ω*-limit sets Ω_1_ and Ω_2_ transitioned into a limit cycle as the frequency of the limit cycle started from non-zero values at the onset of a bifurcation (Fig.2C1), causing discontinuous jumps in the spectrum distance.

**Figure 2.**
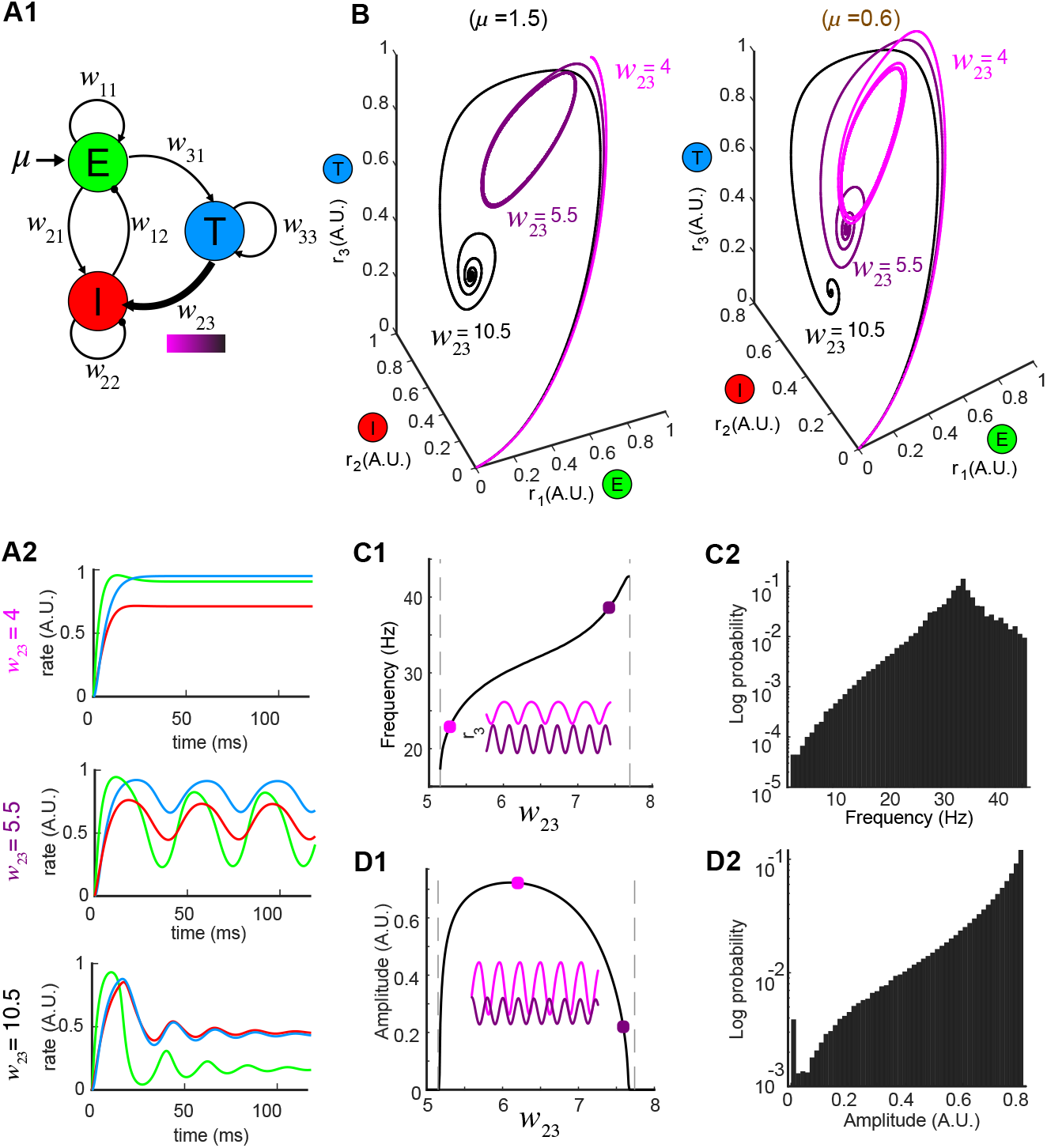
Network dynamics are controlled by weight *w*_23_ of top-down inputs to inhibitory neurons. (A). Changing top-down input *w*_23_ (A1, thick arrow) reshapes network activity to the same stimulus (color bar indicates values of *w*_23_); A2: firing rates at three representative values of *w*_23_ while the stimulus is held constant (*μ* = 1.5). (B). Trajectories in the phase space for the same *w*_23_ as in (A); left: oscillations occur at *w*_23_ = 5.5 for *μ* = 1.5 (same as (A)); right: oscillations occur at *w*_23_ = 10.5 for a different stimulus *μ* = 0.6, illustrating that the dynamics are diverse across different combinations of stimuli and top-down input. (C). Modulation of oscillation frequency by top-down input (for *μ* = 1.5). C1: dependence of frequency on top-down input *w*_23_. Inset: time series of r_3_(t) for two example values of *w*_23_ (squares). C2: a distribution of oscillation frequencies that can be generated by the network for all possible combinations of stimulus and top-down weight. (D). Similar to (C) but for oscillation amplitudes.

However, for the purpose of pattern separation, the two metrics did not give qualitatively different results when assessing the distances due to changes the feedback weight *w*_23_ (Fig.S4, Supplementary materials). Additionally, we found that the non-monotonic dependence of distance in both *d^E^* and *d^S^* on the feedback weight (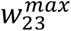) persisted, and that an optimal value for any given pair of stimuli could be found (Fig.S5, Supplementary materials).

### Bifurcation analysis

We denoted the dynamical system of Eqn.(1) as a parameterized form

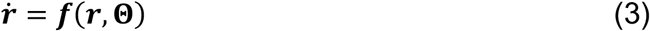

where 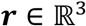 was the vector of firing rates and 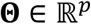 was the vector of parameters. The vector field ***f*** = (*f*_1_, *f*_2_, *f*_3_)^T^ was a smooth function on some open set of 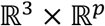. The dimensionality of **Θ** could be up to eight dimensions maximally to include all connection weights *w_ij_* and the external stimulus *μ*. However, since we were only interested in the top-down control, **Θ** was restricted to be two dimensions (*p* = 2) including the top-down input *w*_23_ and the external stimulus *μ*. All the other parameters were fixed as constants determined based on previous experimental work (Whittington, Traub et al. 2000).

As the system Eqn.(3) had an equilibrium at (***r*,Θ**) = (***r***_0_, **Θ**_0_), i.e.,

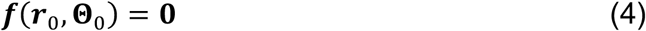

the stability of this equilibrium could be determined from the linearized vector field of Eqn.(3) given by

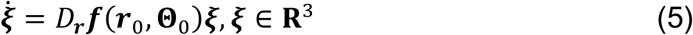

where 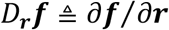 was the Jacobian matrix of the vector field ***f***.

If none of the eigenvalues of *D**_r_f***(***r***_0_, **Θ**_0_) lied on the imaginary axis (i.e., the equilibrium was *hyperbolic*), the local stability of (***r***_0_,**Θ**_0_) in the nonlinear system (4) could be determined by the linear system (5). The equilibrium was stable if all eigenvalues of *D**_r_f***(***r***_0_, **Θ**_0_) had negative real parts. In the case of a hyperbolic equilibrium, varying slightly the parameter **Θ** would not change the stability as taking Eqn.(4) and the invertibility of *D**_r_f***(***r***_0_, **Θ**_0_), there existed a unique smooth function 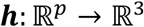 such that

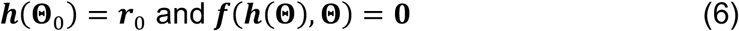

for **Θ** sufficiently close to **Θ**_0_. By continuity of the eigenvalues with respect to parameters, *D**_r_f***(***f***(**Θ**), **Θ**) had no eigenvalue on the imaginary axis for **Θ** sufficiently close to **Θ**_0_. Therefore, the hyperbolicity of the equilibrium persisted and its stability type remained unchanged for in close vicinity of **Θ**_0_.By contrast, when some of the eigenvalues of *D**_r_f***(***r***_0_, **Θ**_0_) lied on the imaginary axis, for example, a zero eigenvalue or a pair of purely imaginary eigenvalues, new topologically different dynamical behaviors occurred by a small change in **Θ**. Equilibria could be created or annihilated, and periodic dynamics could emerge.

The parameterized system (3) thus underwent a bifurcation at (***r***_0_, **Θ**_0_) if the Jacobian matrix *D**_r_f***(***r***_0_,**Θ**_0_) has an eigenvalue of zero real part. In our model, a saddle-node bifurcation (SN) occurred when *D**_r_f***(***r***_0_,**Θ**_0_) had a single zero eigenvalue (in addition to some nondegenerate conditions), and a Hopf bifurcation (H) occurred when *D**_r_f***(***r***_0_, **Θ**_0_) had a pair of purely imaginary eigenvalues. The bifurcation point was found numerically by XPPAUT or the Matlab toolbox MATCONT.

The number of parameters that must be varied simultaneously to evoke a bifurcation is defined as the *codimension* of this bifurcation (Guckenheimer and Holmes 2013, Kuznetsov 2013). Considering the infinite-dimensional space 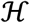 of all vector fields defined on the *n*-dimensional Euclidean space 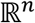, a vector field ***f***_0_ undergoing a bifurcation (Supplemental materials, Fig. S6-S7), for example, a Hopf bifurcation, corresponds to a point in the space 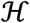. All nearby vector fields with the same singularity as *f*_0_ (i.e., vector fields that are orbitally topologically equivalent to ***f***_0_) form a submanifold 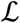 of co-dimension *k*, which is an equivalence class of the singular vector field ***f***_0_. Therefore, within the space 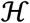 it requires another submanifold 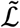 of at least *k* dimensions to intersect transversely with 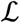 at point ***f***_0_, such that the singularity of ***f***_0_ persists under small perturbations of the vector field. The submanifold 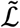 was obtained through a parametrized family of vector fields involving at least *k* parameters. The least number *k* is then defined as the codimension of ***f***_0_. The parametrized vector field ***f***(***r*, Θ**) in Eqn.(3) can be thought of as one realization of the submanifold 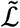 which passes through the vector field 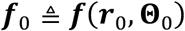 undergoing a bifurcation with the two parameters corresponding to the top-down input *w*_23_ and the external stimulus *μ*.

## Results

### Reduced network model generates complex dynamics

To understand the functional role of top-down projections onto inhibitory neurons, we built a three-node network model (Fig.1A, see Methods) that recapitulates a circuit architecture identified both structurally (Padmanabhan, Osakada et al. 2018) and functionally (Boyd, Sturgill et al. 2012, Markopoulos, Rokni et al. 2012) across a number of brain areas. For different stimuli *μ*, the network exhibited a variety of dynamics (Fig.1B, Fig. S1). For instance, when the stimulus was small, the firing rates *r_i_, i* = 1,2,3 had a fast-transient increase followed by damping oscillations that converged to a stationary state (Fig.1B, top). A sufficiently large stimulus *μ* elevated the firing rates to near saturation, where they then remained at the upper bound of the nonlinear sigmoid function throughout the duration of the stimulus (Fig.1B, bottom). For small or large stimuli, the network responses converged to a constant firing rate after transient dynamics. By contrast, for medium values of *μ*, more complex firing rate dynamics emerged, including oscillations (middle in Fig.1B). To visualize the collective behaviors of E, I, and T populations to these different stimuli, we turned to a three dimensional dynamical system representation of the model where the time evolution of the firing rates (i.e., *state variables*) was as a *trajectory* (or an *orbit*) in the *phase space* (*r*_1_, *r*_2_, *r*_3_) and the tangent vector defining the velocity of each point along a trajectory given by the *vector field **f*** = (*f*_l_, *f*_2_, *f*_3_)^T^ (see Methods) of Eqn.(1). The firing rates over time in Fig.1B thus corresponded to trajectories in Fig.1D starting from the origin 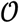 (where all three populations were silent). For small stimuli, the trajectory made an excursion before spiraling into an *equilibrium* indicated by the solid dot (Fig.1D, orange). Similarly, when the stimulus *μ* was large, the trajectory again settled into an equilibrium, but one that was translated within the phase space to the top-right corner (Fig.1D, black). Finally, medium stimuli, the time-varying oscillation of the firing rates manifested as a *periodic orbit* (or a *limit cycle*) in the 3D phase space (Fig.1D, brown). In this space, we defined the steady-state dynamics as the *ω-limit set* of the system.

### Top-down input reshapes network dynamics and modulates neural oscillations

To explore how top-down down projections onto the inhibitory neuron population (I) shaped the dynamics of the network, we studied the effects of changes in the connection weight *w*_23_ on firing rate dynamics. A number of experimental studies have shown that inhibitory neuron activity can be dynamically gated, on short time-scales corresponding to changes in behaviors or internal states such as attention or arousal (Saalmann, Pigarev et al. 2007, Batista-Brito, Zagha et al. 2018), or on longer timescales corresponding to learning and memory (Tomita, Ohbayashi et al. 1999, Cohen, Wilmes et al. 2011). To model these general phenomenological changes, we varied the weight *w*_23_ (Fig.2A, top) from the top-down population (T) to the inhibitory population (I) and studied the effect of these changes on the firing rate dynamics of the network.

For a fixed stimulus (*μ* = 1.5) the dynamics of the firing rates *r_i_*(*t*), *i* = 1,2,3 were sensitive to different values of *w*_23_ (Fig.2A, bottom). When the top down weight was small (*w*_23_ = 4), firing rates approached the equilibrium exponentially (Fig.2A bottom and Fig.2B left, black traces). Conversely, when the top-down weight was large (*w*_23_ = 10.5), the firing rate of excitatory cells (*r*_1_) increased initially, but was suppressed as inhibition reduced the activity, until the firing rates ultimately settled to an equilibrium (Fig.2A top and Fig.2B left, light magenta traces). When the magnitude of top-down input was changed to an intermediate value (*w*_23_ = 5.5), the same stimulus generated oscillatory activity in the network, with the steady-state dynamics transitioning to a periodic orbit (a limit cycle). Changing the weight of top-down projections onto the local inhibitory population (I) in a single stimulus produced the same diversity of firing rate dynamics that occurred from changes in the stimulus. Furthermore, for a given top-down input weight (*w*_23_), the effects on the network dynamics for a particular stimulus was unique to that stimulus (Fig.2B right vs. left).

In regimes where specific weights of top-down input generated sustained oscillatory activity for a given stimulus *μ*, we determined the frequency and amplitude of these oscillations (Fig.2B, left *w*_23_ = 5.5, right *w*_23_ = 10.5) as changes in both have been tied to circuit function and behavior (Buzsaki and Draguhn 2004, Kay, Beshel et al. 2009). For a given stimulus, oscillations emerged between two values of *w*_23_, with the frequency of the oscillation varying monotonically (Fig.2C1). By contrast, while the amplitudes of the oscillations started from zero at the two critical values of *w*_23_, they reached a maximum in between (Fig.2D1). Thus, by changing the top-down input (*w*_23_) the oscillations generated covered a wide range of frequencies (Fig.2C2, 0 – 45 Hz) and amplitudes (Fig.2D2, 0 – 1 A.U.), including frequencies in the alpha, beta and gamma bands (Fig. S2).

### Top-down input contributes to pattern separation

As changing the top-down weight onto inhibitory neurons could generate complex activity patterns we next asked what computations could be performed by this control. For example, both behavioral and neurophysiological measures show that as the representations of two stimuli by neuronal circuits become different, distinguishing between them becomes easier (Leutgeb, Leutgeb et al. 2007, Yassa and Stark 2011). Examples include distinguishing two similar odors that differ slightly in a physiochemical feature (Friedrich and Laurent 2001), or distinguishing between two similarly oriented bars of light (Long, Jiang et al. 2015). In the olfactory bulb, local inhibitory granule cells appear critical for discrimination tasks (Abraham, Egger et al. 2010, Nunez-Parra, Maurer et al. 2013); in cortical circuits, local parvalbumin positive inhibitory neurons play an analogous critical role in discrimination (Korotkova, Fuchs et al. 2010, Lovett-Barron and Losonczy 2014). Both olfactory bulb granule cells and cortical parvalbumin interneurons receive feedback inputs from higher processing areas, suggesting that control of inhibition, via top-down centrifugal projections, may be one way that stimulus discrimination is implemented by the circuit.

To test this hypothesis, we presented our network with a pair of stimuli, denoted by *μ*_1_. and *μ*_2_ (corresponding to stimuli arranged along a one-dimensional axis) and studied how top-down control of inhibition altered the firing rate representations of the two stimuli (Fig.3A). For a set of stimuli *μ_i_, i* = 1,2, the steady-state *representation* of the network activity was the *ω*-limit set Ω*_i_, i* = 1,2 for the associated stimulus with the distance between the two stimuli *μ*_1_ and *μ*_2_ Δ*μ* in the stimulus space, and the resultant distance in the firing rate phase space between the two *ω*-limit sets (Ω. and Ω_2_) as a metric *d* (see Methods). The smaller the Δ*μ*, the more similar the two stimuli were. Thus, for any given pair of stimuli separated by Δ*μ*, we hypothesized that changes in the weight of feedback onto the inhibitory neuron population (*w*_23_) could increase the value of *d*. Such increases in distances would result in the representations of the two stimuli *μ*_1_ and *μ*_2_ becoming more distinct (Fig.3A).

**Figure 3.**
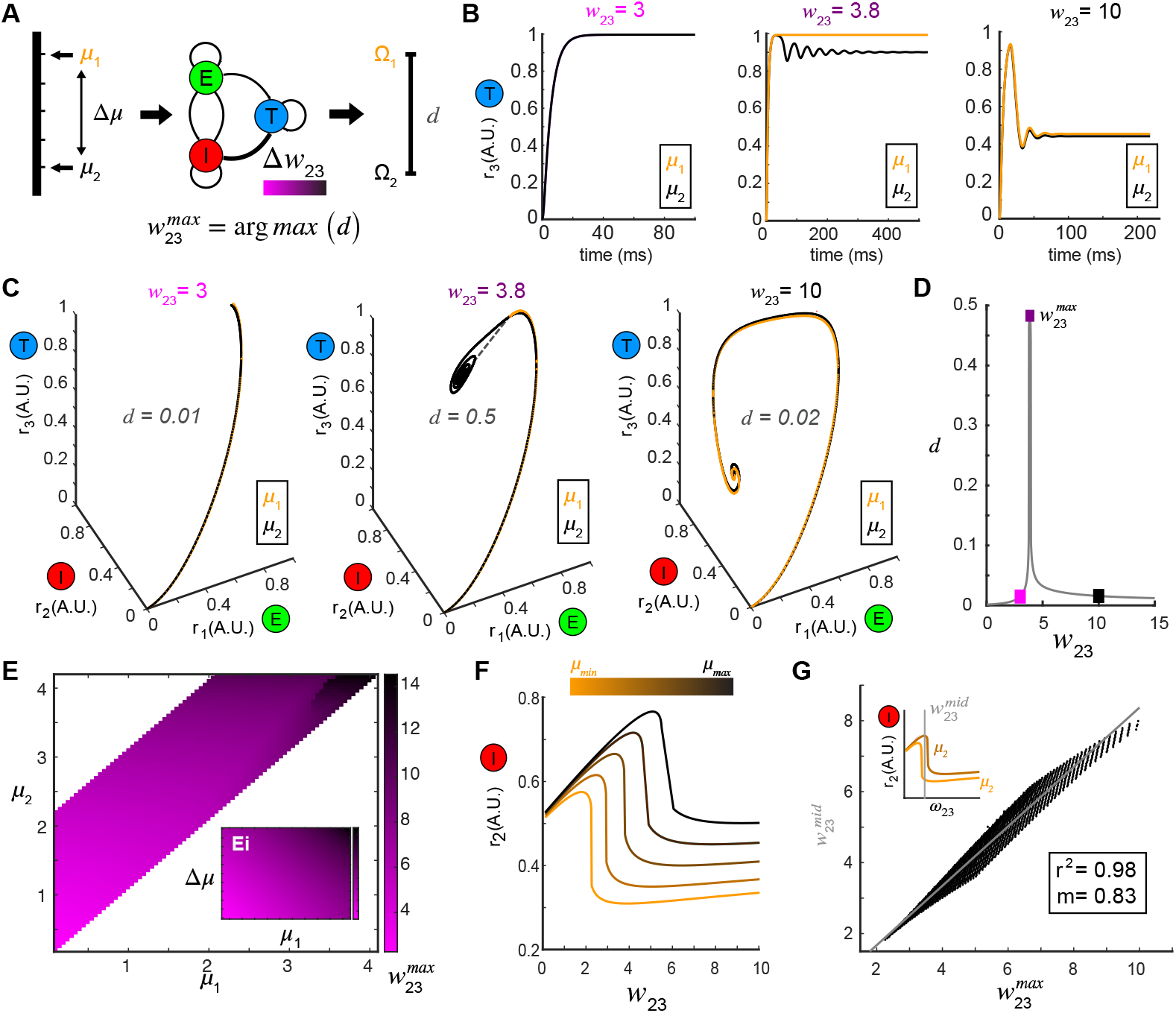
Pattern separation via top-down control of inhibition. (A). A schematic diagram illustrating the separation of response patterns to a pair of stimuli (*μ*_1_ and *μ*_2_) by changing top-down input *w*_23_ (color bar indicates values of *w*_23_). (B). Time series of *r*_3_(*t*) in response to *μ*_1_ = 1.0 and *μ*_2_ = 1.1 at three representative values of *w*_23_ shows that *r*_3_(*t*) are close at some top-down weights (left: *w*_23_ = 3; right: *w*_23_ = 10), but pushed apart at other top-down weights (middle: *w*_23_ = 3.8). (C). Response trajectories and network representations of two stimuli in the phase space from which the distance *d* between representations was calculated. From left to right the top-down input *w*_23_ correspond to those in (B) for the same stimulus pair (*μ*, = 1.0 and *μ*_2_ = 1.1). (D). Nonmonotonic dependence of the distance *d* on top-down weight *w*_23_ (same parameters as in (B)), with maximum achieved at *w*_23_ = 3.8. The squares are color coded as in (B) and (C). (E). The matrix 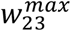 maximizes *d* between network representations in response to all combinations of stimuli (*μ*. and *μ*_2_) presented to the network. Inset: the same matrix of 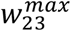 organized by one stimulus *μ*, vs. stimulus difference Δ*μ*. (F). Dependence of the stationary firing rate of inhibitory population *r*_2_ on feedback *w*_23_ at different levels of stimulus (indicated by color bar). (G). The correlation between 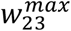 obtained from (E) for all pairs of stimuli *μ*_1_, and *μ*_2_ and the 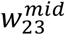 corresponding to the midpoint of the two inhibitory firing rate maxima associated with the same pair of stimuli (upper left inset) reveals that top-down input optimizes pattern separation by gating the balance between excitation-inhibition and recurrent-inhibition. The grey line denotes the utility line. Lower right inset shows the correlation coefficient and the slope of the linear regression.

In a representative example where *μ*_1_ = 1.0 and *μ*_2_ = 1.1, when the top-down input was low (*w*_23_ = 3), the representations of the two stimuli were close (Fig.3B-3C left, *d* = 0.01). As *w*_23_ was increased, the adjacent equilibria were pushed apart (*w*_23_=3.8, Fig. 3B-3C, middle, *d* = 0.5), separating the patterns of activity that corresponded to the two stimuli. Interestingly, as *w*_23_ was increased further (*w*_23_ = 10), the representations of the two stimuli became close to one another again (Fig.3B-3C, right, *d=0.02*). Consequently, the distance *d* between the two equilibria (Ω_1_ and Ω_2_) was a non-monotonic function of *w*_23_ (Fig.3D, Fig. S3). For any given pair of stimuli an optimal value of 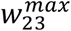 could be found which maximized the distance between the two resultant representations Ω_1_ and Ω_2_. A landscape of the optimal 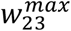 across all pairs of stimuli (*μ*_1_, *μ*_2_) was shown in Fig.3E (Supplemental materials, Fig. S4). Changing the amount of top-down projections onto the inhibitory neuron population (I) could be used to facilitate stimulus separation dynamically (Fig. S5). To understand why top-down inputs achieved this optimal 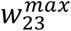, we examined the effect that varying the top-down projection weight had on the stationary responses of both local excitatory and inhibitory populations (*r*_1_ and *r*_2_). For a given stimulus, an increase in *w*_23_ led to a monotonic decrease in *r*_1_, suggesting persistent suppression onto the local E population. By contrast, the response of the local I population *r*_2_ was elevated first with increasing *w*_23_ until reaching the maximum 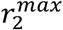, after which *r*_2_ dropped significantly (Fig.3F). Across different levels of stimuli *μ*, the shape of the firing rate *r*_2_ as a function of *w*_23_ remained the same but shifted vertically (Fig.3F). To determine if these differences in the firing rate of inhibitory neurons (*r*_2_) were related to the optimal values of top-down inputs, for any given pair of stimuli (*μ*_1_, *μ*_2_), we plotted the 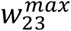 (abscissa) obtained from the landscape in Fig.3E versus the mid-point 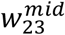 (ordinate) of the connecting line between the 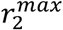 of the same stimulus pair (inset of Fig.3G). The response *r_2_* to one stimulus on the left of the midpoint dropped significantly, while the response to a similar stimulus on the right still had a high inhibitory firing rate. The optimal 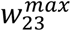 for separating the responses of two similar stimuli was correlated to 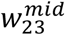 (R^2^ = 0.98), the value at which inhibitory neuron activity from one stimulus was suppressed while activity from the other similar stimulus remained persistently high (Fig.3G). Consequently, stimulus separation arose from the differential sensitivity of inhibitory neurons to the balance between top-down feedback and recurrent inhibition, i.e., when an imbalance occurred between the top-down feedback and the recurrent inhibition for one stimulus while that balance was preserved for the second stimulus.

### Top-down input contributes to oscillation synchrony

In addition to equilibrium attractors in the firing rate phase space across different stimuli, our model was also capable of generating sustained oscillations of firing rates, due to presence of limit cycles. Stimulus-evoked oscillations were modulated by the top-down inputs (Fig.2) and covered a wide range of frequency and amplitude, suggesting the possibility that these oscillatory responses to different stimuli could be synchronized by tuning *w*_23_. To explore this question, we first examined the oscillations in the firing rate generated by two different stimuli *μ*_1_ and *μ*_2_. At a given value of top-down input (*w*_23_ = 12.0), one stimulus (*μ*_1_ = 1.15) generated oscillations (*f*_1_ = 33.5Hz, Fig.4A1, before) in the firing rates of the top-down (T) population that were different in both frequency and amplitude from the oscillations (*f*_2_ = 37.5Hz, Fig.4A1, before) in response to a second stimulus (*μ*_2_ = 0.6). However, a change in the weight of the top-down input (*w*_23_ = 8), resulted in the frequencies of the firing rate oscillations becoming more similar for the same two stimuli (Fig.4A1, after, *f*_1_ = 30.4Hz, *f*_2_ = 30.8Hz). This increase in the firing rate synchrony was also apparent when visualized in the 3D phase space (Fig.4A2). To quantify the synchrony between the two oscillations in response to *μ*_1_ and *μ*_2_, before and after changes in top-down input, we calculated the spectrum distance *d^s^* (see Supplementary material), a measure of the total difference in both frequency and amplitude between the two limit cycles (Fig.4B1). Changing the top-down input resulted in synchronized activity in the network for two different stimuli (Fig.4B2). While the effect was greatest when stimuli were similar, we found examples where oscillations from two different stimuli that were initially as far as 20 Hz apart could nonetheless become synchronous by changing the weight of feedback *w*_23_ onto inhibitory neuron populations. As with stimulus discrimination, a systematic relationship emerged corresponding to the optimal top-down input 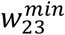 across combinations of stimuli (*μ*_1_ vs. *μ*_2_) that minimized the distance *d^S^* (Fig.4C).

**Figure 4.**
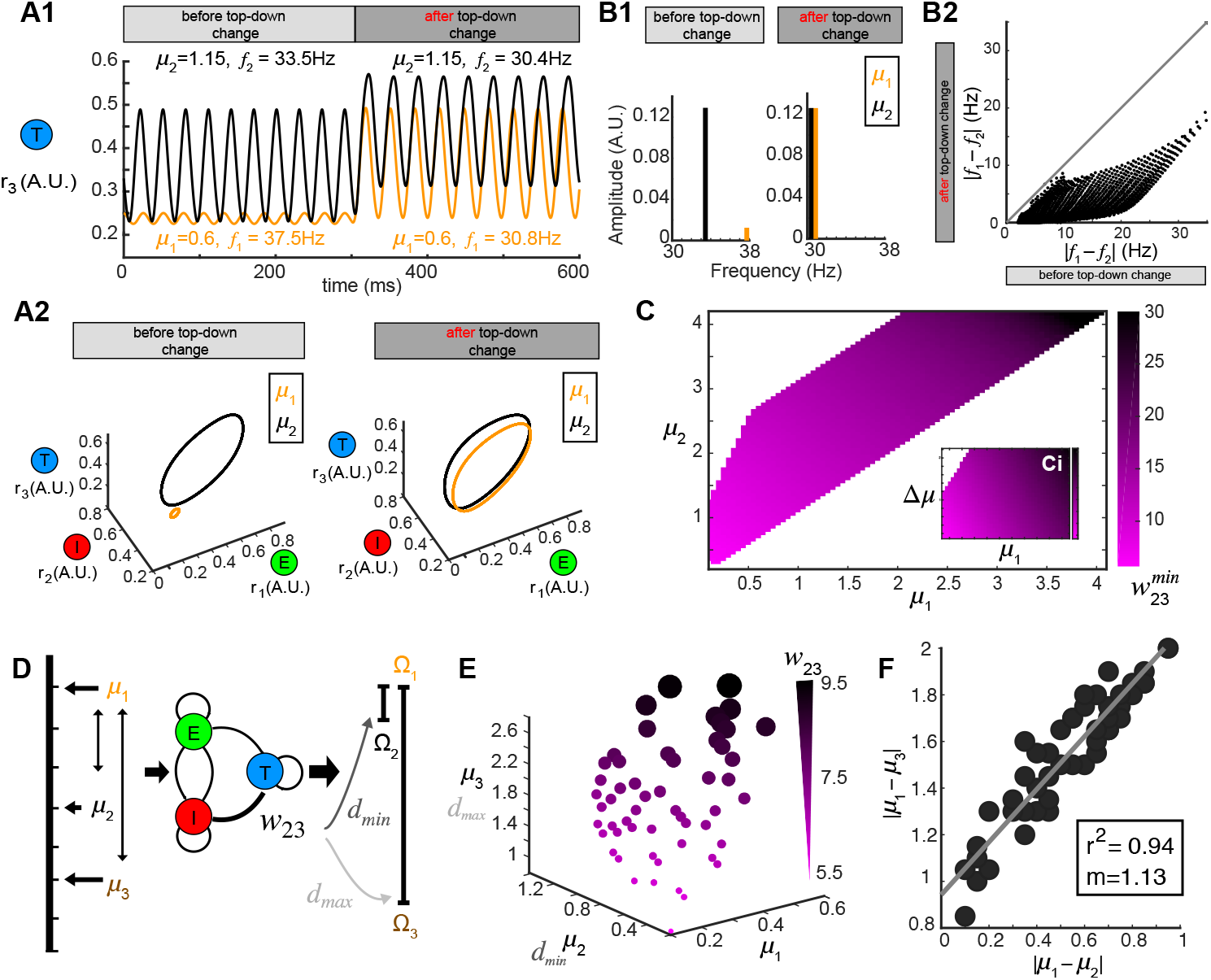
Oscillation synchrony via top-down control. (A). Oscillatory responses to two example stimuli *μ*_1_ = 1.15 and *μ*_2_ = 0.6 become synchronized after changing the top-down input w_23_. A1. Time series of *r*_3_(*t*) before and after changing the top-down input. A2. Limit cycles corresponding to the oscillatory responses in A1 are plotted in phase space. (B). Changing *w*_23_ can make both the frequency and amplitude of two oscillations similar to one another. B1. Frequency and amplitude components of the two oscillations shown in (A). B2. Changing top-down input reduces the frequency differences of responses to two distinct stimuli. (C). The matrix 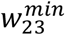 which minimizes the distance *d* between oscillatory responses to all combinations of stimuli *μ*_1_ and *μ*_2_. Inset: the same matrix of 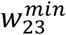 organized by one stimulus *μ*_1_, vs. stimulus difference Δ*μ*. (D). Schematic diagram illustrating that the same value of *w*_23_ which minimizes the distance between oscillations responding to stimuli *μ*_1_, and *μ*_2_ can maximize the distance between responses to stimuli *μ*_1_ and *μ*_3_. (E). Scatter plot in *μ*_1_ - *μ*_2_ - *μ*_3_ space where each sphere size and color corresponds a top-down weight *w*_23_. (F). Correlation between the differences of stimuli whose response distances are simultaneously minimized and maximized. Inset: correlation coefficient and the slope of linear regression.

Although we thus far treated stimulus discrimination and response synchrony separately, neural circuits perform both operations simultaneously, bringing the network representation of one stimulus closer to another, while simultaneously pushing the representation of that stimulus farther from a third. We therefore tested if a single change in the top-down input *w*_23_ accomplish both of these operations; minimize the distance between the responses to one pair of stimuli (*μ*_1_ vs. *μ*_2_) while also maximize the response distance to another other pair of stimuli (*μ*_1_ vs. *μ*_3_, Fig.4D). To do this, we identified values of *w*_23_ that were optimal for both synchrony between the oscillations generated by stimulus *μ*_1_ and μ_2_ (Fig.4C) and also produced a maximum separation between the representations of stimulus *μ*_1_ and *μ*_3_. A 3D scatter plot of *μ*_1_ – *μ*_2_ – *μ*_3_ (Fig.4D), showed all examples corresponding to two pairs of stimuli: (*μ*_1_, *μ*_2_) whose oscillatory responses to stimuli were synchronous for a given *w*_23_ and two stimuli (*μ*_1_, *μ*_3_) whose responses were also maximally separable at the same *w*_23_. The values of top-down input *w*_23_ for each point that fulfilled these diametrically distinct functions were coded by color and size (Fig.4E). Importantly, we found that the top-down input scaled with the stimuli, such that when *μ*_1_, *μ*_2_ and *μ*_3_ were small, the top-down input that achieved both synchrony and separation was also small, but as the three stimuli increased in magnitude, the top-down input needed to synchronize one pair and separate the other pair also increased, suggesting that the optimal weights for the top-down projections onto inhibitory neurons that performed both pattern separation and oscillation synchrony corresponded to the magnitude of the stimuli. Finally, we found a strong correlation between the values of stimulus differences: |*μ*_1_ – *μ*_2_| and |*μ*_1_ – *μ*_3_| (Fig.4F) at which an *w*_23_ weight was optimal for stimulus separation and oscillatory synchrony.

### Bifurcation mechanism for top-down control of inhibition

Finally, to understand mathematically how such operations emerged as a result of changes in the top-down weight to inhibition, we studied the structure of the transitions in the dynamics of firing rates in the network (Fig.2A). These transitions were associated with qualitative or topological changes in the *ω*-limit sets of the system, indicative of the occurrence of bifurcations in the system. To explore this further, we first examined the how the *ω*-limit sets of the system changed with different top-down inputs. An equilibrium corresponding to constant firing rates in the network (Fig.5A, left and right) arose from the intersection of three nullclines (Fig.5A, yellow thick lines), each characterizing the geometric shape on which the firing rate derivatives of two nodes equaled to zero (Fig.S6, Supplementary materials). Global phase structures for two representative values of top-down inputs (Fig.5A, left: *ω*_23_ = 15, right: *ω*_23_ = 4) illustrated how these equilibria varied within the firing rate phase space, where the purple arrows showed the velocities (directions and magnitudes) of the flows along which trajectories followed. Using this information, for a given combination of stimulus *μ* and top-down input *w*_23_, we calculated a family of *ω*-limit sets on which dynamics settled from any set of initial values in the firing rates of the excitatory (E), inhibitory (I), and top-down (T) populations. In these two examples, both equilibria were stable and attractive, with all nearby trajectories (black thin lines) moving towards them. This was, however, not true for all values of *w*_23_. At some critical values of *w*_23_, the equilibrium lost stability, and a small-amplitude limit cycle branched from that unstable equilibrium, resulting in the oscillations observed in the dynamics (Fig.5A, middle). This transition signified a *Hopf* bifurcation of the system (see Methods and Supplementary materials, Fig. S6), which arose when the top-down input *w*_23_ was tuned within a specific regime. Across combinations of external stimuli *μ* and top-down input *w*_23_, we obtained a smooth manifold in the phase space (Fig.5B, left), composed of stable and unstable equilibria and divided into several regions. Sustained oscillations corresponded to the red region-LC (LC: limit cycle) where each equilibrium (unstable) was paired with exactly one limit cycle that were born simultaneously via a Hopf bifurcation (the purple empty square vs. the dot-dashed curve). Constant firing rates corresponded to the grey region-EE (EE: exponential equilibrium) and green region-SE (SE: spiraling equilibrium), where the equilibria were stable, approached either exponentially (region-EE) or via damping-oscillations (region-SE). Finally, the two blue regions on the manifold were bounded by another type of bifurcation near Bogdanov–Takens (BT) and an example global phase structure (Fig.S7). The equilibrium manifold thus defined the entire family of network representations (i.e., the of *ω*-limit sets) for all possible combinations of stimuli *μ* and top-down input *w*_23_.

**Figure 5.**
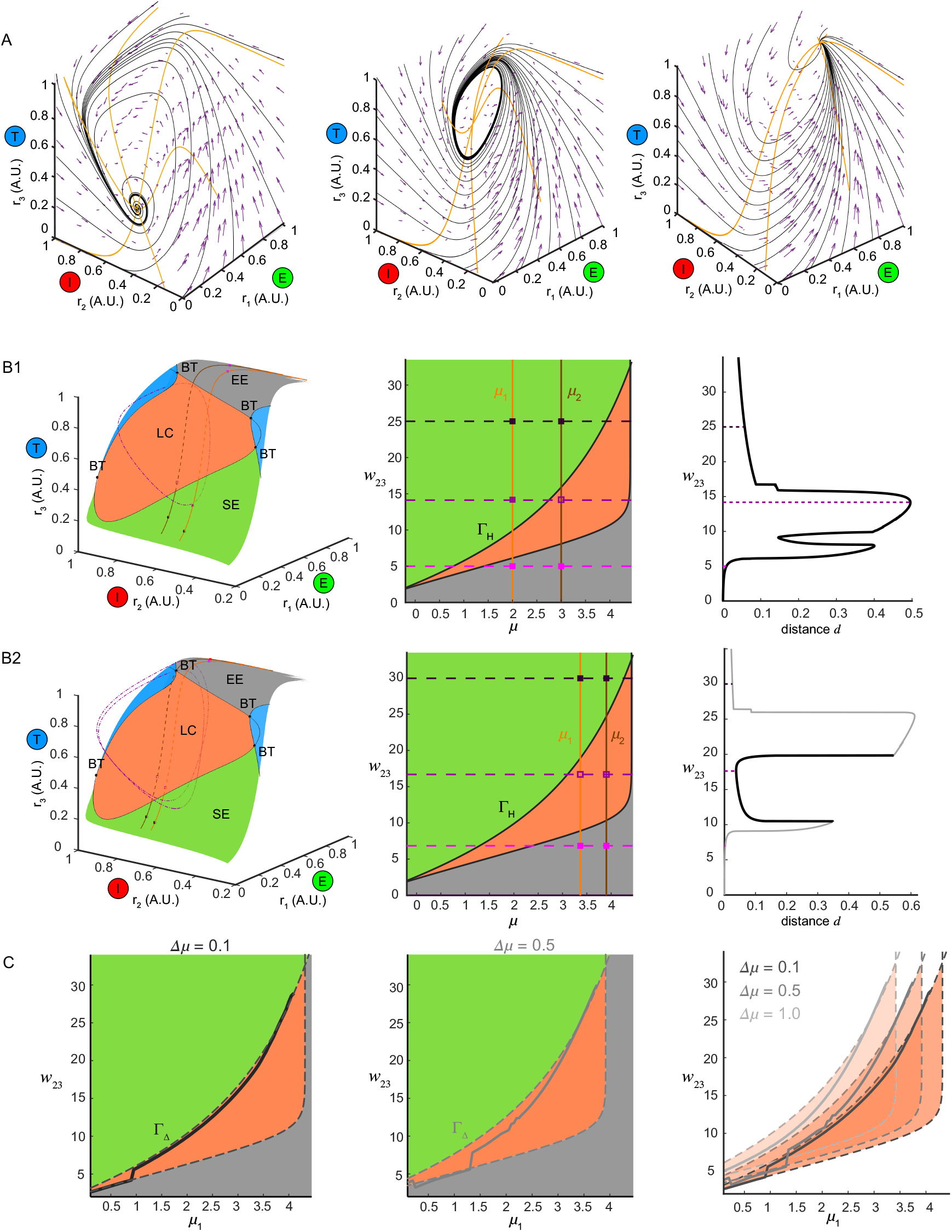
Bifurcation mechanism of top-down control to support both stimulus separation and oscillation synchrony. (A). Global phase structures showing the vector field (purple arrows), nullclines (yellow thick curves) and several representative trajectories (black thin curves) for the same stimulus *μ* = 1.5 and three different values of *w*_23_. Left, *w*_23_ = 15; middle, *w*_23_ = 6; right, *w*_23_ = 4. (B). Illustration of the bifurcation mechanism and the transition boundary of dynamics in phase space and parameter space. B1, pattern separation. Left: the equilibrium manifold in the firing rate phase space divided by the separatrix from multiple codimension-2 bifurcation points: BT (Bogdanov–Takens bifurcation) into several regions. Region-LC: limit cycle. Region-SE: spiraling equilibria. Region-EE: exponential equilibria. Blue regions: regions where multiple equilibria coexisted.. For two example stimuli *μ*_1_ = 2.0 and *μ*_2_ = 3.0 given in the middle of B2, two paths of equilibria were induced on the equilibrium manifold and traversed across different regions as changing top-down input *w*_23_. Middle. Different regions on the equilibrium manifold corresponded to different regimes in the parameter space of *μ* and *w*_23_ in the same color scheme (parameters for blue regions in B1 were largely beyond the range in B2 thus not shown). The transition boundary Γ_H_ specified the pair (*μ*,*w*_23_) at which the network underwent a Hopf bifurcation and corresponded to the separatrix enclosing the region-LC in B1. Two given stimuli were denoted by two vertical lines and three example values of *w*_23_ corresponded to three horizontal dashed lines, giving rise to a pair of junctions for each. These junctions were also plotted as squares in the left of B1 denoting the corresponding *ω*-limit sets in the same color (solid square: stable equilibrium; empty square: unstable equilibrium). Right: the distance between the *ω*-limit sets to represent the two given stimuli. The maximal distance was achieved when the two junctions were on opposite sides of Γ_H_. B2: same as B1 but for oscillation synchrony occurring when the two junctions were both inside Γ_H_. (C). Comparisons between the translated transition boundary Γ_Δ_ (dashed curve) depending on Δ*μ* (left: Δ*μ* = 0.1, middle: Δ*μ* = 0.5) and the sliced section of the 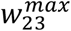 at the same Δ*μ* (solid curve). Right: a series of translated Γ_Δ_ for three representative values of Δ*μ*.

Within the space of *μ* and *w*_23_, we identified a transition boundary Γ_H_ (black solid curves, Fig.5B, middle) corresponding to the separatrix enclosing the region-LC on the equilibrium manifold. Γ_H_ specified the parameter pairs (*μ, w*_23_) at which a Hopf bifurcation occurred, thereby dividing the parameter space into regimes where different dynamics were present (same color coded as Fig.5B1, left). For a given pair of stimuli (for example, *μ*_1_ = 2.0, *μ*_2_ = 3.0), changing *w*_23_ corresponded to shifting the horizontal dashed line vertically (three representatives were shown in Fig.5B1, middle), thereby shifting the junctions with the two stimuli (vertical solid lines, Fig.5B1, middle) across different regimes in the parameter space. Correspondingly, in the firing rate phase space (Fig.5B, left), changes in *w*_23_ for one stimulus moved the equilibrium through different regions of the manifold: EE-LC-SE (the part in region-LC was plotted in a dashed curve), while for another stimulus, a parallel curve on the manifold could also be traced by changing *w*_23_. When the two junctions in Fig.5B1 (middle) were on opposite sides of the transition boundary Γ_H_, with one equilibrium in region-LC and the other in region-SE (Fig.5B1, left), the two *ω*-limit sets (i.e., network representations) became topologically different from each other; the former a limit cycle, and the latter an equilibrium point. For this specific combination of stimulus pairs and top-down input, the distance between the network representations of the two stimuli became maximal (Fig,5B1, right) such that the optimal 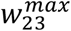 for pattern separation was then achieved when the *ω*-limit sets were on different sides of transition boundary Γ_H_.

Furthermore, when the two junctions were both inside the transition boundary Γ_H_ (Fig.5B2, middle) for a pair of stimuli, this time, when the top-down input was altered for both, two limit cycles emerged (one for each stimulus, for example, *μ*_1_ = 3.45, *μ*_2_ = 3.95, Fig,5B2). As both the frequencies and amplitudes of the oscillations were modulated by *w*_23_ (Fig.2C and 2D) and synchrony between the network responses for the two stimuli were achieved when the top-down input was tuned to place both junctions within this space. As changes in top-down input moved the junctions for pairs of stimuli within the parameter space, a shared mechanism supported both stimulus separation and oscillation synchrony,depending on the relative positions of the junctions with respect to the transition boundary.

We next wished to determine if the transition boundary Γ_H_ identified via analysis of the dynamical system corresponded to the 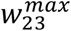 matrix found in Fig.3Ei, a measure of the computational optimization performed by the network. To do this, we considered a set of *μ*_1_, i.e., {*μ*_1_| *μ*_1_ ∈ [0,4]}, and a set of Δ*μ*, wherein each value was an array of stimuli *μ*_1_ + Δ*μ* that defined a unique set of stimulus pairs {(*μ*_1_, *μ*_1_ + Δ*μ*)| *μ*_1_ ∈ [0,4]}. For a given Δ*μ* > 0 (distinguishing the pair (*μ*_1_, *μ*_1_ + Δ*μ*) was the same task as distinguishing (*μ*_1_ + Δ*μ, μ*_1_) in terms of a pattern separation), for instance, Δ*μ* = 0.1 or Δ*μ* = 0.5 (Fig.5C, left and middle), the set of stimulus pairs had a unique transition boundary Γ_Δ_ (Fig.5C, right). The section of the 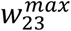 matrix in Fig.3Ei for the set of stimulus pairs {(*μ*_1_, *μ*_1_ + Δ*μ*)| *μ*_1_ ∈ [0,4], Δ*μ* is given} was correlated to the transition boundary Γ_Δ_ of the same value Δ*μ* (Fig.5C). For small Δ*μ* = 0.1, the slice of the 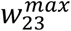 followed closely with the transition boundary Γ_Δ_ (Fig.5C, left). As the stimulus difference increased (Fig.5C, middle), 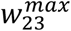 deviated from the boundary Γ_Δ_. Larger Δ*μ*’s increased the bifurcation lag between two stimuli such that the one that bifurcated first had more parameter space to develop before the bifurcation of the other stimulus. This suggested that as the stimulus discrimination became harder, i.e., as Δ*μ* decreased, the range of top-down input *w*_23_ to separate two stimuli shrank significantly around a close vicinity of the transition boundary. Consequently, accurate and subtle adjustments of top-down input around the transition boundary were then required to separate similar stimuli from each other. Taken together, we identified a bridge linking the mechanisms that gave rise to the dynamics of this neural circuit with the computations that were performed by the circuit.

## Discussion

Using a three-node model, which includes top-down centrifugal projections onto inhibitory neurons, we identified a network capable of complex dynamic behaviors, ranging from an equilibrium attractor to stable oscillations across a range of frequencies and amplitudes. By changing the weight of top-down projections onto inhibitory neurons, the network could either facilitate pattern separation across two similar stimuli, or synchronize the oscillatory activity produced by two different stimuli. A bifurcation analysis of the dynamical system revealed that both mechanisms emerged from the transition boundary of Hopf bifurcations which branched from co-dimensional two bifurcation points (i.e., the Bogdanov-Takens bifurcation). Our results provide both a mathematical framework for how top-down control of inhibition confers unique properties to the dynamics of networks, and a link as to how these dynamics support the computations the neural circuits are often performing.

An important point is to consider how these changes in top-down projections may be instantiated biologically? The mechanisms governing such changes are dependent on the timescale of weight changes.

First, evidence suggests that behavioral state can strongly modulate the local dynamics of a circuit, including the firing rates of inhibitory neurons (Pakan, Lowe et al. 2016, Hattori, Kuchibhotla et al. 2017). For example, in primate v4, inhibitory neurons show strong attentional modulation (Fries, Reynolds et al. 2001, Mitchell, Sundberg et al. 2007). Similarly, synchronization of parvalbumin (PV) inhibitory neurons and elevated gamma has been tied to attention (Kim, Ährlund-Richter et al. 2016). Additionally, it has been suggested that neuromodulatory signals may gate the effect of attention (Thiele and Bellgrove 2018). Changes in the amount of top-down feedback that occur on short timescales such as during attention, may thus be implement by neuromodulators that affect either the release of neurotransmitter, or gate the responses at the post-synaptic target of glutamate release. Either would change the weight of top-down inputs onto inhibition dynamically. Both dopamine (Gao and Goldman-Rakic 2003, Tritsch and Sabatini 2012) and serotonin (Athilingam, Ben-Shalom et al. 2017) alter excitatory post-synaptic potential (EPSP) amplitudes in inhibitory interneurons, and such alterations would result in changes in the weight of synapses from top-down projections to local inhibitory interneurons. In this way, either pattern separation, or oscillatory synchrony could happen on relatively short timescales corresponding to epochs of attention. Here the modulation of top-down weight would be flexible and dynamic.

Second, top-down weights could change on longer timescales as a result of synaptic plasticity (Isaacson and Scanziani 2011, Kullmann, Moreau et al. 2012). While initial reports identified plasticity at glutamatergic synapses on inhibitory interneurons (Buzsaki and Eidelberg 1982, Tomasulo and Steward 1996), the rules governing this plasticity appear to be more complex than the rules for glutamatergic plasticity on excitatory neuron synapses. Nonetheless, classic plasticity induction paradigms, such as pairing theta burst stimulation with post-synaptic depolarization (Perez, Morin et al. 2001, Lamsa, Irvine et al. 2007), result in changes in the weight of glutamatergic synapses on inhibitory interneurons. In this example, top-down changes in the weight would correspond to long timescales, operating to codify stable changes in synapses that render representations permanently more distinct or similar.

While the biological mechanisms by which the weights of top-down synapses change onto inhibitory neuronal populations may be diverse depending on the timescale of that change, they can give rise to functionally equivalent changes to the dynamics of neural circuits that support an array of computations. For instance, changes in the top-down weight to inhibitory neurons would render two stimuli more distinct at the level of firing rates in the population, a process referred to as pattern separation (Cayco-Gajic and Silver 2019) or decorrelation (Friedrich and Laurent 2001). Our model predicts that pattern separation may arise as a feature of the non-monotonic change in firing when the weight of feedback is gradually altered. The amount of top-down projections onto inhibitory neurons thus sets a gate, allowing some stimuli to cross the threshold of inhibition, while others do not; the consequence of which is increasing their distance in representational space.

In parallel, changing top-down weights onto inhibitory neurons could increase the oscillatory synchrony in the network for two stimuli that were initially asynchronous. A number of experimental and theoretical studies have explored the privileged role that inhibitory interneurons play in generating gamma oscillations (Whittington, Traub et al. 1995, Hasenstaub, Shu et al. 2005, Cardin, Carlén et al. 2009, Sohal, Zhang et al. 2009, Tiesinga and Sejnowski 2009). Among these, the two most common models are those where gamma arises from reciprocal coupling between pyramidal cells and inhibitory interneurons (PING), and recurrent connections among inhibitory interneurons (ING) (Whittington, Traub et al. 2000, Tiesinga and Sejnowski 2009). In both, the generation of oscillatory activity arises as a result of motifs of local connectivity. In our work, we identified another motif by which gamma oscillations arise – **T**op-down control of **I**nhibitory **N**euron **G**amma (TING). Local excitatory cells broadcast activity patterns to a top-down population, that then synapses onto a local inhibitory interneuron population. As the system crosses the transition boundary defined by a combination of stimulus and top-down weight, gamma oscillations emerge. In addition to identifying how gamma oscillations can emerge in a circuit with top-down connections made exclusively onto local inhibitory interneuron populations, we also demonstrate how synchronous oscillatory activity within local circuits can be changed by changing the weight of top-down projections onto inhibition.

Finally, our model links two key areas of computational neuroscience, the dynamical systems framework for evaluating how neuronal responses evolve over time, and principles of neural coding, the aim of which is to understand what can be done computationally with the temporal evolution of neural dynamics. The capacity of the top-down feedback to modulate not only the dynamics of the local network, but to reshape its own input information via inhibition reveals an essential feature of neural circuits. Using bifurcation analysis, we found that the system was governed by a codimension-2 bifurcation, with distinct domains within a manifold of the system defined by the stimulus and the weight of feedback. These domains corresponded to transitions in the dynamics of the system. Changes in the top-down inputs onto inhibition operated by moving a transition boundary for dynamics across different external stimuli. When two stimuli were on opposites sides of this transition boundary, their dynamics operated under two different regimes, and their representations were pushed further apart. By contrast, when the stimuli were both on the same side of the transition boundary, within regimes corresponding to similar dynamics, their activity became more synchronous; effectively binding those stimuli together. Changes in top-down weight were therefore changes in the location of the transition boundary that could either marshal the representations of two stimuli together or push them apart.

In conclusion, we identified a model that links the dynamics of neural systems with the computations they are hypothesized to perform, and may be used as a generalized framework to study the diverse effects of feedback onto inhibitory populations.

## Acknowledgments

This study was supported by funding from the National Institutes of Health (NIH) and the National Science Foundation (NSF). KP was funded by NIH R01 MH113924, NSF CAREER 1749772, the Cystinosis Research Foundation, and the Kilian J. and Caroline F. Schmitt Foundation.

## Supplementary materials

### Stimulus of intermediate strength evoked oscillations of firing rates

Our network model generated diverse dynamics in response to different stimuli *μ*. As the input increased from low to high values, the steady states of the dynamics transitioned from equilibria to limit cycles and then back to equilibria again (Fig. 1). We plotted firing rates of the top-down population (T) after transient dynamics as a function of input magnitude (Fig S1). An intermediate interval existed where *r*_3_ occupied a continuum of firing rates for one stimulus, indicating the emergence of oscillations. However, when the stimulus strength was beyond that interval, either too low or too high, the firing rate *r*_3_ achieved a constant value for the stimulus.

### Feedback modulation of neural oscillations across different feedforward levels

For an input of intermediate value (*μ* = 1.5) changing the connection weight *w_23_* corresponding to the top-down projection to inhibitory neurons modulated the oscillations over a range of frequencies as well as amplitudes (Fig. 2). The control of both the frequency and amplitude via changes in *w*_23_ occurred across an array of weights (*w*_31_) associated the feedforward drive from the local excitatory population (E) to the feedback population (Fig.S2). Interestingly, the range of *w*_23_ within which neural oscillations emerged became narrower with increasing *w*_31_, suggesting that the feedforward drive established the dynamic range within which changes in feedback (*w*_23_) adjusted both the amplitude and frequency of the oscillations that could be generated by the network.

### Definition of period of a limit cycle

From the perspective of dynamical system, a limit cycle in the phase space corresponds to the oscillation of firing rates in the temporal space. Since the time constants in our model had units of millisecond, the frequency of oscillations was defined as 1000/*T*, with T denoting the period of limit cycles, which was defined as follows: if *r_i_*(*t*), *i* = 1,2,3 denote the firing rate of neural population in the model, then a limit cycle satisfies the periodicity:

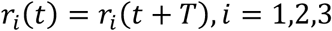

for some T>0 and all 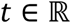. The minimal T for which the above equality holds is the period of the limit cycle.

### Definition of Euclidean distance *d^E^* and spectrum distance *d^s^*

Here we defined two types of distances, the Euclidean distance *d^E^* and the spectrum distance *d^s^*, to measure the separation between the *ω*-limit sets for different stimuli. The Euclidean distance was defined as follows: supposing that Ω_1_ and Ω_2_ are two *ω*-limit sets composed of *N*_1_ and *N*_2_ discrete points in three-dimensional phase space (*r*_1_, *r*_2_, *r*_3_) respectively, denoted as {***α***_1_, ***α***_2_,…,***α***_*N*_1__} and {***β***_1_, *β*_2_,…, ***β***_*N*_2__} where *α_i_* and *β_j_* are three-dimensional vectors, then

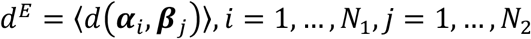

Where 〈·〉 denotes the average and 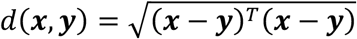 which is the standard distance between any two points in the three-dimensional Euclidean space. Note that in the case of equilibria, the number of discrete points in the ω-limit set was one: *N* = 1, and in the case of a limit cycle, we set *N* = T/*dt*, where *T* was the period and *dt* was time bin for numerical integration.

The Euclidean distance *d^E^* worked well in measuring the distance between two equilibria in the phase space, but when one ω-limit set was a limit cycle, the averaging operation in the definition made it only a coarse and lagged estimate of the separation for equilibrium vs. limit cycle and limit cycle vs. limit cycle. In particular, the activity of the top-down population (*t*) should be decodable with respect to the stimulus information, something that was problematic for when using *d^E^*. Therefore, we defined the spectrum distance *d^S^* to address the question of distances between representations that were sensitive to those representations being oscillations. To calculate the spectrum distance *d^S^* between two ω-limit sets Ω_1_ and Ω_2_, only the sequence of their *r*_3_ component was decomposed by Fourier transform which converted the firing rate signal in the temporal domain into a representation in the frequency domain.

The single-sided amplitude spectrum for the Fourier transform of the firing rate signal *r*_3_(*t*), was used to obtain peaks around frequency values. For an equilibrium corresponding to constant firing rate *r_3_* = *A*, there existed only one peak around zero frequency with its amplitude proportional to *A*, since the Fourier transform of a constant function is a delta-function. We referred this component as the direct component (DC) of the signal. For the case of a limit cycle, in addition to one peak around zero frequency, there existed another peak around frequency 1000/*T* where *T* denoted the period of the limit cycle. We referred this additional peak of a limit cycle as the alternating component (AC). Thus, the spectrum distance *d^S^* was a sum of the differences between both direct components (DC) of two limit sets and alternating components (AC) of two limit sets, which was formalized as follows: supposing that *D*_1_ and *D*_2_ were the amplitudes of the peaks at zero frequency for two *ω*-limit sets Ω_1_ and Ω_2_, and ***a**_i_* = (*f_i_,A_i_*), *i* = 1,2 denoted the corresponding alternating components of Ω_*i*_, where was the non-zero frequency and *A_i_* was the amplitude of the peak around *f_i_*, then we have

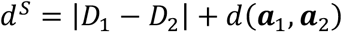

Note that we set ***a***_*i*_ = (0,0) if the *ω*-limit sets Ω_*i*_ was an equilibrium. Thus, when the two limit sets were both equilibria, *d^S^* only contained the first term measuring the difference between the direct components. In this case, *d^S^* was only a linear projection (up to a constant factor) of the Euclidean distance *d^E^*.

### Landscape of distance *d^E^* and *d^S^* over different input pairs

The network model generated an *ω*-limit set in response to each input, which was taken as the network representation of the external stimulus. Using the distance defined above, we measured the separation between two network representations given any pair of stimuli: *μ*_1_ and *μ*_2_ = *μ*_1_ + Δ*μ*. Although the absolute value of *d^E^* between a given stimulus pair was different from the value of *d^S^* between the same pair, the overall trends of the landscapes over all combinations of stimuli given by *d^E^* and *d^S^* were found to be similar (Fig.S4). When the feedback was small (*w*_23_ = 1.5), the landscapes of *d^E^* and d^s^ varied smoothly across all stimulus combinations. This was because the only form of *ω*-limit sets in this case was an equilibrium. However, a relatively larger feedback weight (*w*_23_ = 4) unmasked complicated waved structures confined within the range of stimuli (*μ*_1_ < 1). Also, the magnitude of both *d^E^* and d^S^ were increased by an order, suggesting the representations reflected larger distances, that would improve the discriminability between two inputs. *d^E^* and *d^S^* consequently quantified different attributes of how one limit set was separated from the other one. The Euclidean distance *d^E^* characterized their relative configurations in the three-dimensional phase space, while the spectrum distance *d^S^* accounted for the oscillatory extent by decomposing the one-dimensional signal into DC and AC. Therefore, *d^S^* was more sensitive to the transitions of the forms of limit sets, as well as the difference of frequencies between two limit cycles. The 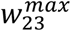 matrices given by the two distances were compared (Fig.S5) and we found that for most input pairs the differences 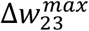 remained small. The one given by the Euclidean distance was shown in Fig.3.

### Global phase structures - nullplanes and nullclines

For some fixed external stimulus (*μ* = 1.5), changing the feedback weight *w*_23_ would change the global phase structures, which in turn gave rise to changes of network dynamics. In our three-dimensional model, three “nullplanes” could be obtained by equaling the right hand side (RHS) of each component of Eqn.(1) to zero, on which the velocity vectors only had two non-zero components (Fig.S6). Furthermore, three “nullclines” were pairwise intersections of nullplanes, on which the velocity vectors only had one non-zero component. Therefore, the equilibrium of the system was given by the intersection of all nullclines, where the velocity was a zero vector. Changing feedback weight *w*_23_ tilted one nullplane (blue) determined by the second equation of Eqn.(1), thus moving the positions of the intersection point, i.e., the equilibrium. Around *w*_23_ = 6 (middle), the equilibrium was unstable and there existed a limit cycle surrounding it; a Hopf bifurcation occurred during this process.

### Conditions for Hopf bifurcation

First, starting from *w*_23_ = 15, the linearized Jacobian matrix of the vector field of Eqn.(1) at the equilibrium had a pair of complex-conjugate eigenvalues with negative real parts (the third eigenvalue is negative real number. In this case nearly all trajectories in the phase space spiraled into this equilibrium referred to as a *focus-node* so that the time sequences exhibit a transient phase of damping oscillations (Fig.2A). Then slightly altering the parameter *w*_23_ caused small variations of the equilibrium position as well as the associated eigenvalues, until some critical point of *w*_23_ when this pair of complex-conjugate eigenvalues crossed the imaginary axis in the complex plane. As the real parts of the complex pairs became zero, the equilibrium lost stability and a small-amplitude limit cycle branched from that unstable equilibrium, i.e., a Hopf bifurcation.

### Bistable regions in the manifold of equilibria

There were two blue regions (referred to as BS) on the manifold of equilibrium in Fig.5B, left, which were bounded by saddle-node bifurcations near BT. By choosing the representative parameters as μ = –1, *w*_23_ = 0.2, we found that the nullclines intersected at three points (Fig.S7A, yellow thick lines), two of which were stable and the other one was a saddle point (with two positive eigenvalues and one negative eigenvalue). Therefore, in this case two stable equilibria coexisted in the phase space and the entire phase space was the divided into two basins of attraction (BOA) by the stable manifold of saddle point and the nullplane (Fig.S7B) corresponding to the second equation of Eqn.(1). Trajectories initiating from within one BOA (Fig.S7A, black thin lines) approached the equilibrium belonging to it.

**Fig.S1.**
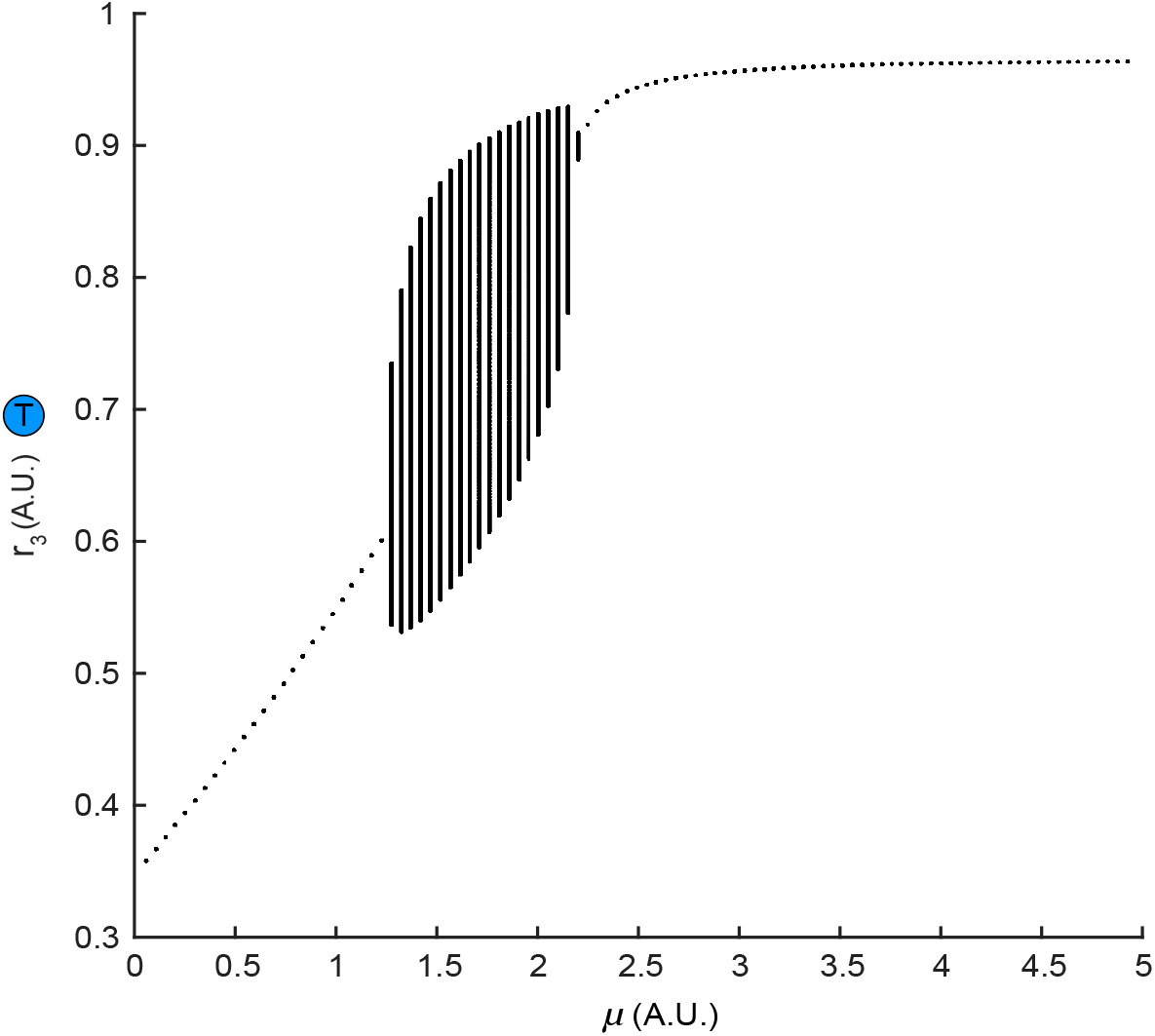
Transitions of dynamics elicited by varying stimulus strength. External stimulus of intermediate strength (approximately *μ* ∈ [1.4,2.5]) evoked sustained oscillations of firing rates. Individual points indicated constant firing rate and continuous vertical lines indicated oscillations.

**Fig.S2.**
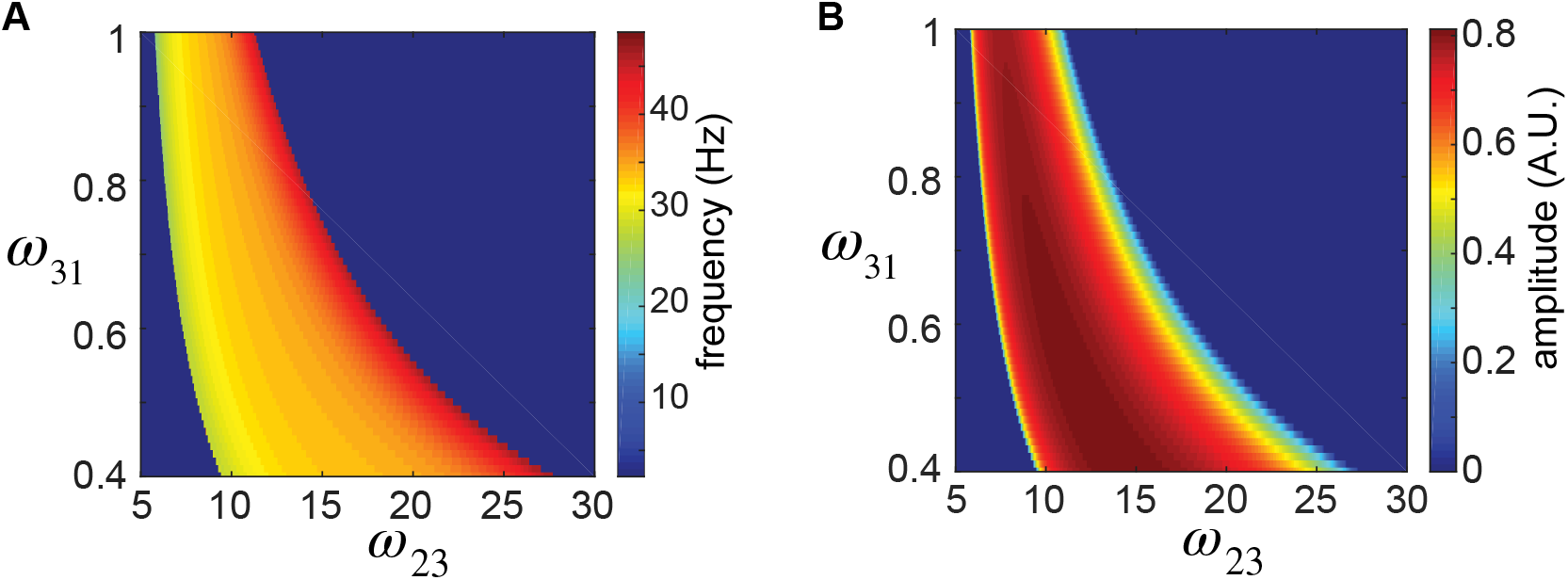
Dependence of oscillation frequency and amplitude on the feedforward and feedback weight (fixing *μ* = 1.5). Oscillations occurred within an interval of feedback weight *w*_23_ whose size shrank as the feedforward weight *w*_23_ increased. (A). Heatmap of frequencies. (B). Heatmap of amplitudes.

**Fig.S3.**
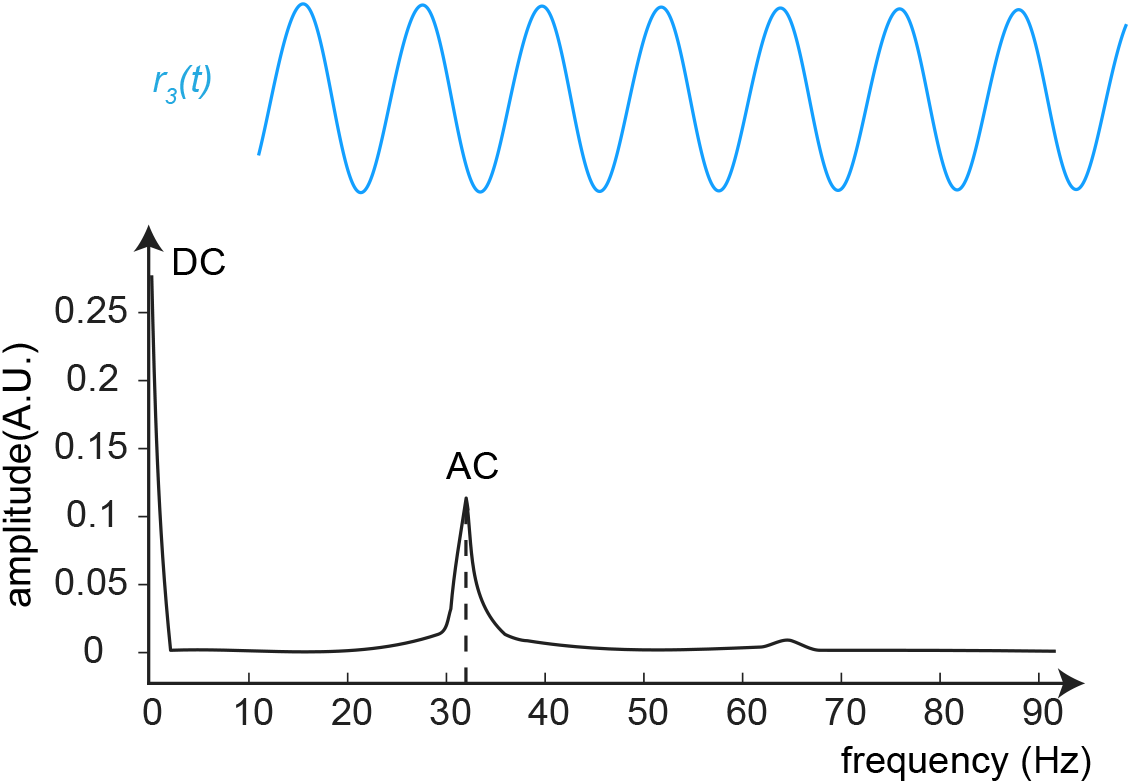
Schematic illustration of direct component (DC) and alternating component (AC) extracted from the time series of *r*_3_(*t*) by Fourier transform. Top: a representative waveform of oscillatory firing rate signal of *r*_3_(*t*). Bottom: the components of the signal above. The direct component (DC) at zero frequency was proportional to the average value of the periodic signal. The alternating component (AC) measured the maximum deviation of firing rate from DC, which gave the amplitude of AC, and how frequent that deviation happened, which gave the frequency of AC.

**Fig.S4.**
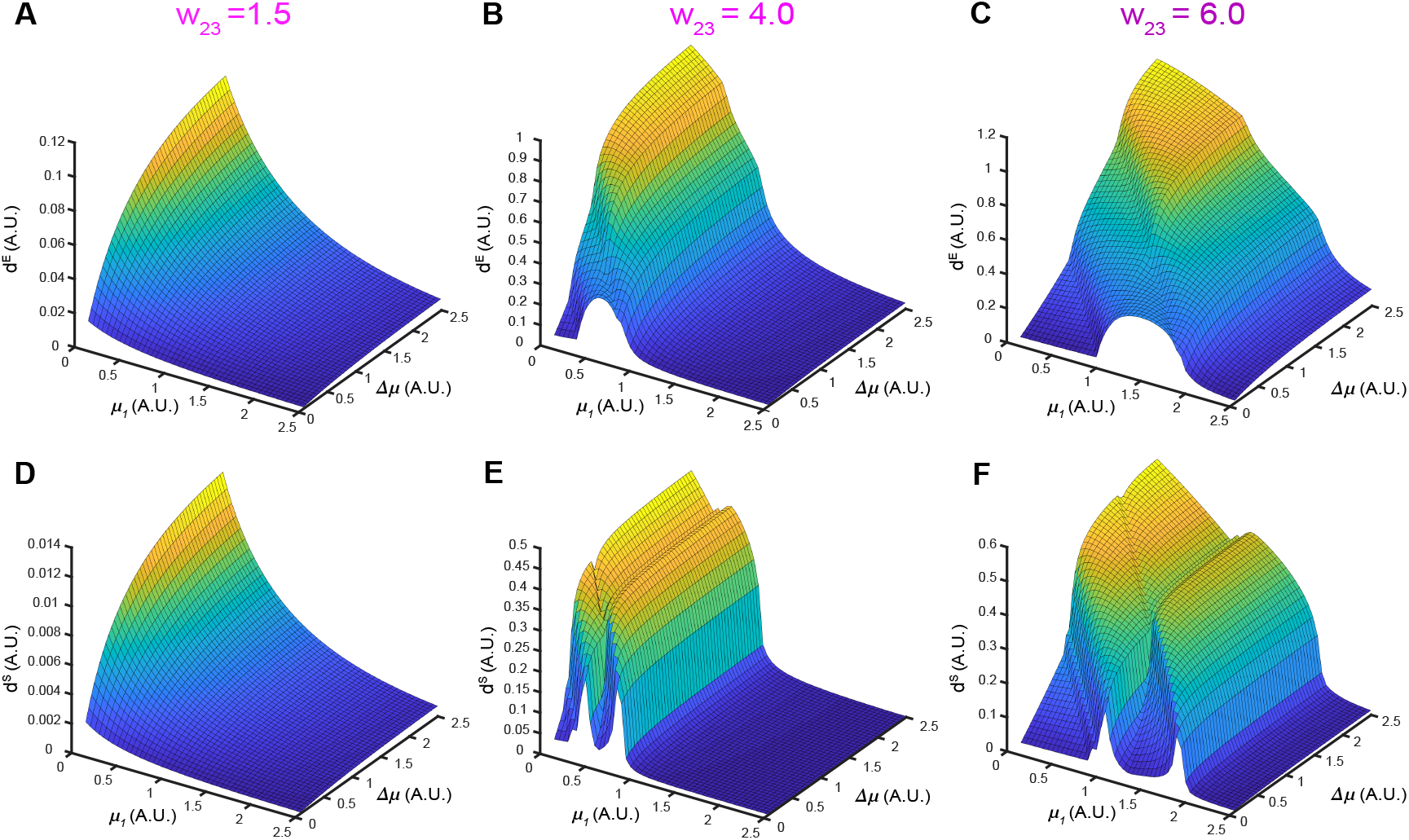
The evolution of distance landscapes between two *ω*-limit sets generated by different pairs of stimuli (*μ*_1_ and *μ*_2_ = *μ*_1_ + Δ*μ*) as the feedback weight *w*_23_ varied. (A-C). Landscapes given by the Euclidean distance for three example values of *w*_23_. (D-F). Landscapes given by the spectrum distance for the same values of *w*_23_ as in (A)-(C).

**Fig.S5.**
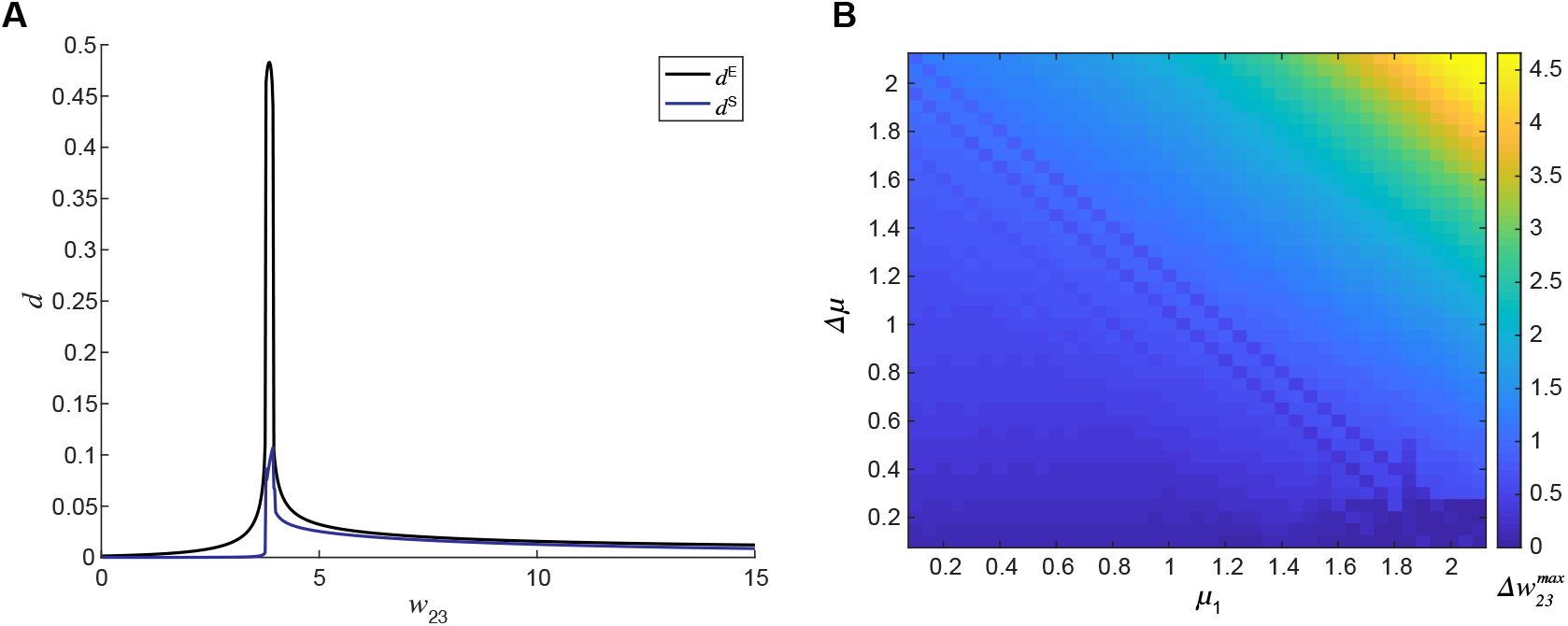
No qualitative difference between results given by the Euclidean distance and spectrum distance. (A). The non-monotonic dependence of both *d^E^* and *d^S^* on the feedback weight *w*_23_ for the same pair of input stimuli (*μ*_1_ = 1.0, *μ*_2_ = 1.1). Both metrics gave similar results of 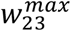. (B). Matrix of the difference in 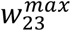 given by *d^E^* (Fig.3) and *d^S^*. The color bar indicated the absolute values of the differences.

**Fig.S6.**
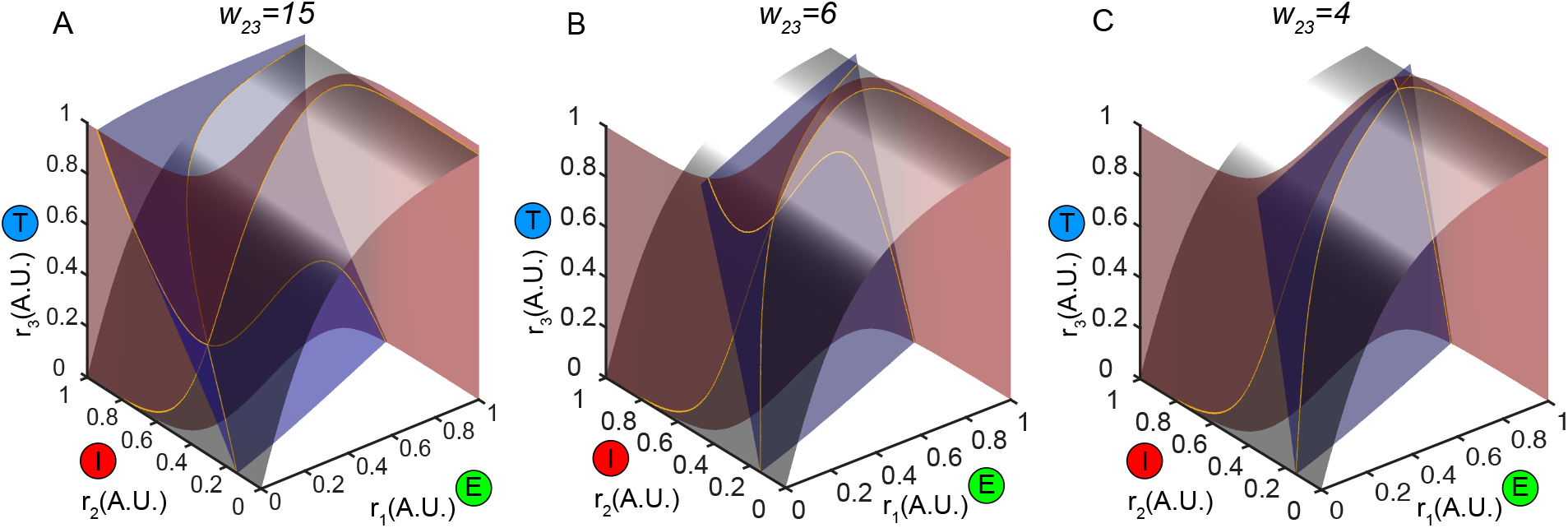
Global phase structures at different example values of *w*_23_. Three nullplanes were denoted as transparent surfaces whose intersections gave rise to nullclines. Changing the feedback weight tilted the blue nullplane progressively to change the position of the equilibrium as well as its stability.

**Fig.S7.**
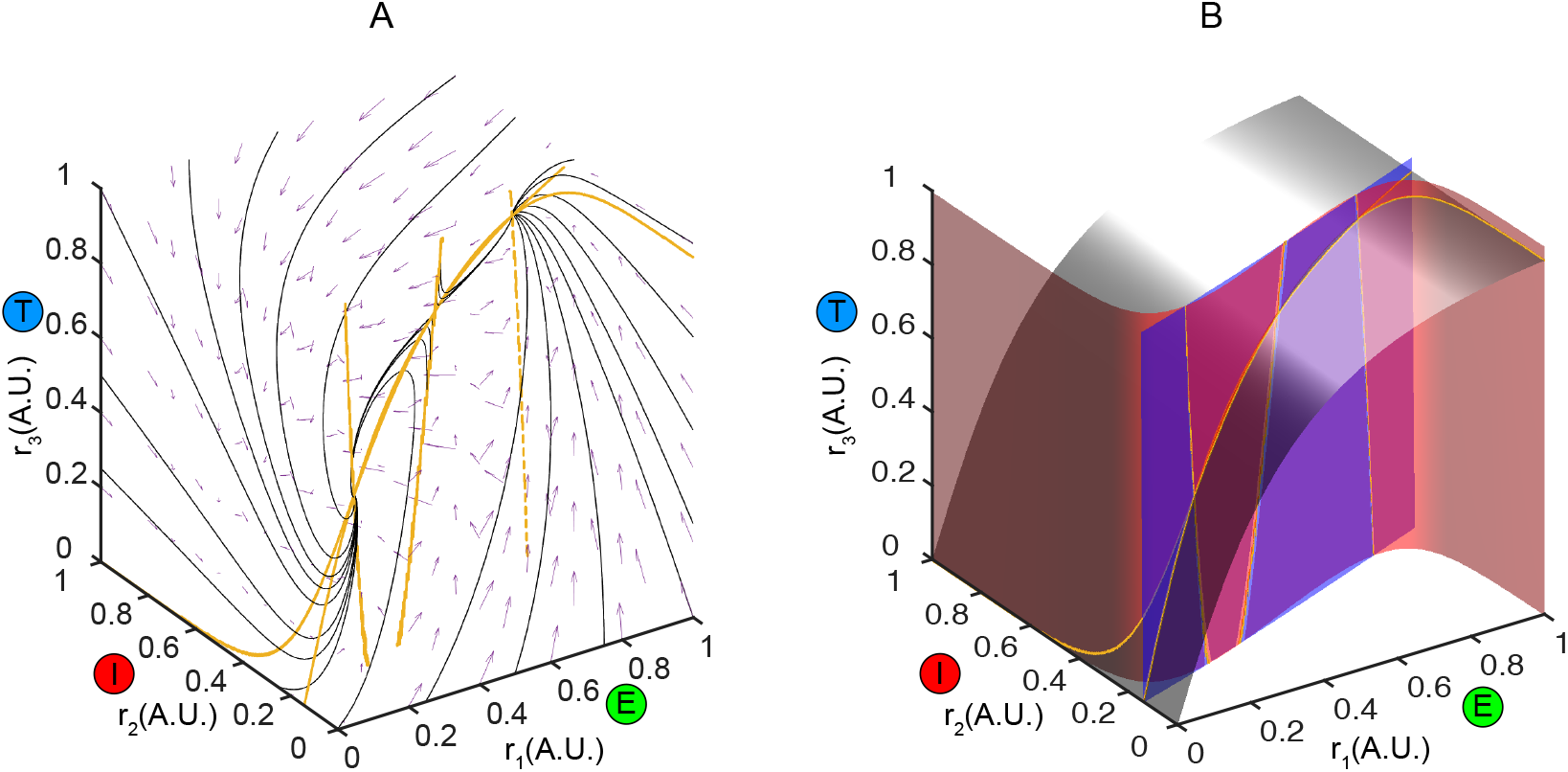
Global phase structures of bistability. (A). Nullclines (yellow thick curves) intersected at three equilibria, two of which were stable and attracting nearby trajectories (black thin curves). The middle equilibrium was a saddle point. Purple arrows indicated the velocities of flow. (B). Nullplanes giving rise to the nullclines in (A).

